# *Comammox* and Unknown Candidate AOB Contribute to Nitrite Accumulation in an Integrated A-B stage process that Incorporates Side-stream EBPR (S2EBPR)

**DOI:** 10.1101/2023.03.29.534650

**Authors:** Yuan Yan, Jangho Lee, IL Han, Zijian Wang, Guangyu Li, Kester McCullough, Stephanie Klaus, Da Kang, DongQi Wang, Anand Patel, Jim McQuarrie, Beverley M. Stinson, Christine deBarbadillo, Paul Dombrowski, Charles Bott, April Z. Gu

## Abstract

A novel integrated pilot-scale A-stage high rate activated sludge, B-stage short-cut biological nitrogen removal and side-stream enhanced biological phosphorus removal (A/B-shortcut N- S2EBPR) process for treating municipal wastewater was demonstrated with the aim to achieve simultaneous and carbon- and energy-efficient N and P removal. In this studied period, an average of 7.62 ± 2.17 mg-N/L nitrite accumulation was achieved through atypical partial nitrification without canonical known NOB out-selection. Network analysis confirms the central hub of microbial community as Nitrospira, which was one to two orders of magnitude higher than canonical aerobic oxidizing bacteria (AOB) in a B-stage nitrification tank. The contribution of comammox Nitrospira as AOB was evidenced by the increased amoB/nxr ratio and higher ammonia oxidation activity. Furthermore, oligotyping analysis of Nitrospira revealed two dominant sub-clusters (microdiveristy) within the Nitrospira. The relative abundance of oligotype II, which is phylogenetically close to Nitrospira_midas_s_31566, exhibited a positive correlation with nitrite accumulation in the same operational period, suggesting its role as comammox Nitrospira. Additionally, the phylogenetic investigation suggested that heterotrophic organisms from the family Comamonadacea and the order Rhodocyclaceae embedding ammonia monooxygenase and hydroxylamine oxidase may function as heterotrophic nitrifiers. This is the first study that elucidated the impact of integrating the S2EBPR on nitrifying populations with implications on short-cut N removal. The unique conditions in the side-stream reactor, such as low ORP, favorable VFA concentrations and composition, seemed to exert different selective forces on nitrifying populations from those in conventional biological nutrient removal processes. The results provide new insights for integrating EBPR with short-cut N removal process for mainstream wastewater treatment.

## 1 Introduction

The adsorption/bio-oxidation (A-B) process is a promising engineering application towards carbon capture, energy recovery, and efficient nutrient removal. It diverts organic carbon to anaerobic digestion and thus manipulates the C/N in B-stage reactors through carbon flow redirection (Ren and Pagilla, 2022). The first stage (A-stage) is designed for high-rate carbon capture and removal using high-rate activated sludge (HRAS), which is widely operated with a low hydraulic retention time (HRT), a low sludge retention time (SRT) and a low dissolved oxygen (DO) concentration less than 0.5 mg L^−1^ (Boehnke et al., 1997). After HRAS, the wastewater COD/N ratio could be lowered to 0.67 (Lotti et al., 2015), which is suitable for second stage (B-stage) nutrient (mostly nitrogen), like short-cut nitrogen removal processes, to achieve “carbon-free” nitrogen removals (Sancho et al., 2019).

Partial nitrification with anammox (PN/A) process has emerged to be one of the promising short- cut N removal system, which can save 100% carbon, 60% aeration and reduce 80% sludge production (Jetten et al., 1997). It consists of two step processes where aerobic ammonia-oxidizing bacteria (AOB) oxidize half of the ammonia to nitrite and, then anaerobic ammonia-oxidizing bacteria (anammox) bacteria convert ammonia and nitrite into dinitrogen. The out-selection of nitrite oxidizing bacteria (NOB) in B-stage is crucial for PN/A process and it has been successfully demonstrated for side-stream high-strength wastewater (Lackner et al., 2014). A number of operational strategies have been suggested to enable side-stream NOB out-selection and consequently nitrite accumulation, including high temperature, low DO, low SRT, high pH, and high free nitrous acid (FNA) and/or free ammonia (FA) concentration (Hellinga et al., 1998; Joss et al., 2009; Van Dongen et al., 2001; Xu et al., 2012). However, these strategies were not readily applicable in mainstream treatment since the inhibitory levels of 0.02 mg/L FNA and 1 mg/L FA is unavailable with only 50 mg-N/L ammonia at neutral pH in the mainstream (Wang et al., 2017). Also, it has been shown that NOB can adapt to high concentration of FA (An et al., 2021) and low DO (Liu and Wang, 2013), which makes it harder to be out-selected in mainstream process. Therefore, maintaining long-term stable NOB out-selection still remains a challenge for main-stream wastewater treatments and the underlined mechanisms need to be explored.

Most PN/A studies focus only on nitrogen removal, however, phosphorus co-occur with nitrogen and is also a critical nutrient that needs to be treated in wastewater (Rittmann et al., 2011). Enhanced biological phosphorus removal (EBPR) process has been widely applied worldwide in full-scale facilities. Its performance and stability strongly rely on the activity of phosphorus accumulating organisms (PAOs) with favorable influent rbCOD/TP ratio >15 (Gu et al., 2008). Therefore, EBPR is considered incompatible with B-stage autotrophic nitrogen removal due to the insufficiency of readily biodegradable carbon in A-stage effluent (B-stage influent). The implementation of a side-stream anaerobic biological sludge hydrolysis and fermentation reactor called side-stream EBPR (S2EBPR) can be an alternative option to address the carbon shortage challenge of coupling EBPR with B-stage nitrogen removal process. S2EBPR takes advantage of the volatile fatty acids (VFAs) generated via hydrolysis/fermentation of side-stream return activated sludge (RAS) and mixed liquor, and thus overcome the reliance on the influent carbon (Barnard et al., 2017; Onnis-Hayden et al., 2020; Gu et al., 2018; Wang et al., 2019). However, the influence of implementing S2EBPR on the PN/A removal system, especially on the nitrite accumulation remains unknown. We hypothesize that the incorporation of S2EBPR would affect the nitrifying populations because of the extended anaerobic conditions in the side- steam reactor with low oxidation reduction potential (ORP <300 mV) and high VFAs. In addition, PAOs and the surplus of sludge fermentate from the S2EBPR reactor to the down- stream B-stage would compete oxygen with nitrifiers (Carvalheira et al., 2014).

Recently, our team reported a successful incorporation of S2EBPR into a A-B stage process to achieve simultaneous nitrogen and phosphorus removal for main-stream treatment (McCullough et al., 2023). This pilot-scale system combined A-stage HRAS, B-stage short-cut biological nitrogen removal process, and S2EBPR, referring as A/B-shortcut N-S2EBPR herein, provides us a good opportunity to explore the interacting impacts of PAOs and nitrogen removal related microorganisms. In this study, we focused on the comprehensive examination of the nitrite accumulation and AOB and NOB population dynamics in B-stage processes that generated nitrite with intermittent aeration AvN (ammonia versus NO_x_^–^-N (nitrite+nitrate) control strategy. Process performance, batch activity tests were performed to investigate the underlying processes involved in the nitrite accumulation. 16S rRNA gene amplification sequencing and quantitative PCR were applied to reveal the microbial community composition and dynamics with emphasis on nitrification-relevant populations. To our knowledge, this is the first study that revealed the influence of implementing S2EBPR on B-stage nitrification population and, provided new insights on the distinctive nitrifying populations that contributed to nitrite accumulation in this A/B-shortcut N-S2EBPR process that is different from those typically observed in previous WWTPs. The results pointed out a possible new strategy in achieving nitritation to enable anammox for main-stream municipal WWTPs.

## 2 Material methods

### 2.1 Pilot reactor setup and Operation conditions

The pilot plant operated by Hampton Roads Sanitation District (HRSD) was located at Chesapeake-Elizabeth WWTP in Virginia Beach, VA, USA. It consisted of an A-stage HRAS process followed by the B-stage process, a parallel S2EBPR reactor and a final stage anammox polishing tank (Figure S1). A portion of A-stage wasted activated sludge (WAS) was split into a combined fermenter and thickener. The B-stage consisted of four CSTRs bioreactors in series, each with independent aeration control via AvN control (Regmi et al., 2014). In the studied period, B-stage reactors were switched from clarifier wasting to aeration tank wasting (hydraulic wasting) and recycling. A portion of B-stage RAS and the A-stage WAS fermentate were added to the side-stream reactor for PAOs selection. The detailed parameters of A-B process have been introduced elsewhere (McCullough et al., 2023).

Chemical analysis of soluble COD, OP, total ammonia nitrogen (NH_4_^+^-N + NH_3_-N), NO_2_^−^ -N, and NO_3_^−^ -N, volatile fatty acids (VFA) including acetic, propionic, butyric, isobutyric, valeric, isovaleric, and caproic acids were summarized in Text S3.

### 2.2 AOB and NOB activity analysis

AOB and NOB activities were analysis weekly on site. About 4L mixed liquor suspended solids (MLSS) were withdrawn from the last B-stage aerobic tank and pre-aerated for 30 min to oxidize the residual carbon. Ammonium chloride was spiked to reach 25 mg-N/L final concentration. DO and pH were maintained around 3-5 mg/L and 7, respectively. The samples were taken every 15 minutes for a total of 1 hour. After filtered by 0.45 μm filter, the filtrate was used to analyze the NH_4_^+^-N, NO_2_^−^ -N, and NO_3_^−^ -N concentrations using HACH TNT kits (HACH Loveland, CO). AOB activity was calculated as the sum of nitrite and nitrate production rates, and NOB activity was calculated as the nitrate production rates by linear regression.

### 2.3 Quantification of nitrifying candidates

Quantitative polymerase chain reaction (qPCR) was performed on a CFX96 real-time PCR detection system (Bio-Rad, USA) to determine the concentrations of functional genes including *amoA*, *nxrB*, *amoB* and 16S. The amoA-1F/2R primer for the canonical ammonia monooxygenase (AMO) enzymes in AOB (Rotthauwe et al., 1997), and the *nxrB* 169F/638R primer for the beta subunit of nitrite oxidoreductase (NXR) (Pester et al., 2014) were used to target canonical AOB and NOB. The *amoB* 148F/485R primer were used to target the beta subunit of AMO in *Nitrospira* that can do complete ammonia oxidation (comammox) (Cotto et al., 2020). All functional genes were then normalized by 16S rRNA with primer set 341f/534r (He et al., 2007). Calibration curves were obtained using 10-fold serial dilutions of known concentrations of cloned DNA samples that were prepared following the instruction from TOPO TA cloning kit (Thermo Fisher, USA) in our laboratory. The R^2^ of standard curves were above 0.99 and the amplification efficiency was around 95%. The detailed qPCR reaction conditions were summarized in Table S1.

### 2.4 Bioinformatics and statistical analysis

#### 2.4.1 Microbial population analysis

Sludge samples from B-stage were monthly collected for DNA extraction. Genomic DNA was extracted using Instagene matrix (Kim et al., 2013) and then used for both 16S ribosomal RNA (16S rRNA) sequencing and functional genes quantification.

The V4 region of the 16S rRNA genes were amplified using primers 515F-Y (Parada et al., 2016) and 926R (Quince et al., 2011). Detailed amplification process is listed in text S2. To minimize the inflation of rare species in the community analysis, we removed the operational taxonomic units (OTUs) that are lower than 0.01%. The filtered-OUT matrix was used to perform statistical analyses in R-4.2.3. The matrix was also used to predict the bacterial metagenomes using Phylogenetic Investigation of Communities by Reconstruction of Unobserved States 2 (PICRUSt2) pipeline (Douglas et al., 2020) with implemented tools EPA-ng (Barbera et al., 2019), gappa (Czech et al., 2020), and castor (Louca and Doebeli, 2018). The network analysis of microbial community and the correlation between the environmental variables and the most abundant 50 taxa were done through microbiomeSeq package (Ssekagiri et al., 2017; Torondel et al., 2016). All bioinformatic figures were generated with ggplot2 package (Villanueva and Chen, 2019, p. 2).

#### 2.4.2 Oligotyping analysis

Oligotyping analysis was performed according to (Eren et al., 2013; Srinivasan et al., 2021) to reveal the sub-clusters (i.e. microdiveristy) within the dominant nitrifying population. A total of 65,24316S rRNA gene amplicon sequences classified as *Nitrospira* were extracted and formatted as required by the oligotyping pipeline. After quality filtering, 18 nucleotide positions were selected for entropy decomposition (102, 120, 134, 135, 149, 151, 160, 181, 197, 199, 210, 219, 240, 262, 265, 375, 488, 491), which were used to generate a total of 205 raw oligotypes. After denoising and elimination based on the -s (oligotypes that occurred in at least 3 samples) and -M (a minimum count of 30 (∼0.1%)) parameters, 2 dominant oligotypes, presumably representing micro-diversity of *Nitrospira* sub-groups, were finally identified. The representative sequence of the 2 dominant oligotypes were BLAST’d against a custom database built from MiDAS 4.8.1 for top hits at 99% identity threshold (Dueholm et al., 2022).

## 3 Results and Discussion

### 3.1 Simultaneous nitrite accumulation and efficient P removal in A/B-shortcut N- S2EBPR process

The pilot was designed to perform nitrogen and phosphorus removal simultaneously. To overcome the shortage of influent VFA and enable bio-P removal in main-stream treatment line, a side-stream reactor was implemented to facilitate bio-P removal through receiving supplemental carbon from A-stage sludge fermenter (Figure S1). With the increase in the A-stage WAS fermentate supplement flow and RAS slit ratio to the S2EBPR reactor (Figure S2), the nitrite accumulation was observed over a period of approximately 150 days (Figure 1). During this period, the effluent nitrite concentration raised to a maximum level of 6.55 mg-N/L with an average concentration of 2.65 ± 1.89 mg-N/L (Figure 2A). The average nitrite accumulating ratio (NAR) was 49%, suggesting that a significant nitrite accumulation occurred. Figure 2A shows the N mass balance in the system and changes of the various N species along the B-stage reactors in series. As ammonia decreased along the B-stage zones, the nitrite began to accumulate from 3.9 g/d to 20.7 g/d (Figure 2A). Previous research indicated that nitrite can inhibit PAO activity by increasing maintenance energy and disturbing the TCA cycle (Zhou et al., 2011). Nevertheless, our results indicated an efficient P removal (77%-92%) could be achieved concurrently with nitrite accumulation (0.2 ± 0.05 - 2.7 ± 1.9 mg-N/L) during this same period (Text S1, Figure S2).

**Figure 1.**
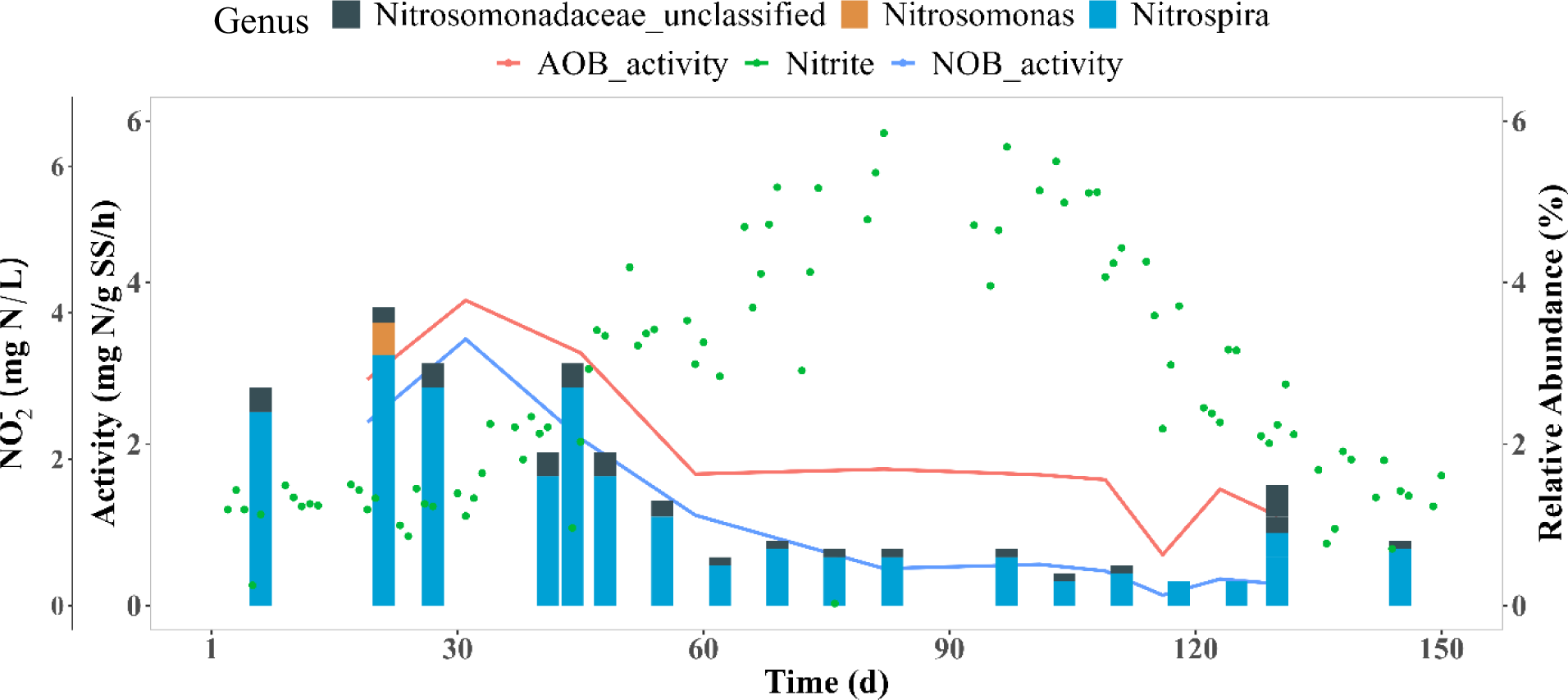
AOB and NOB activities, nitrite accumulation, and the relative abundance of AOB and NOB throughout the operation period for this study.

**Figure 2.**
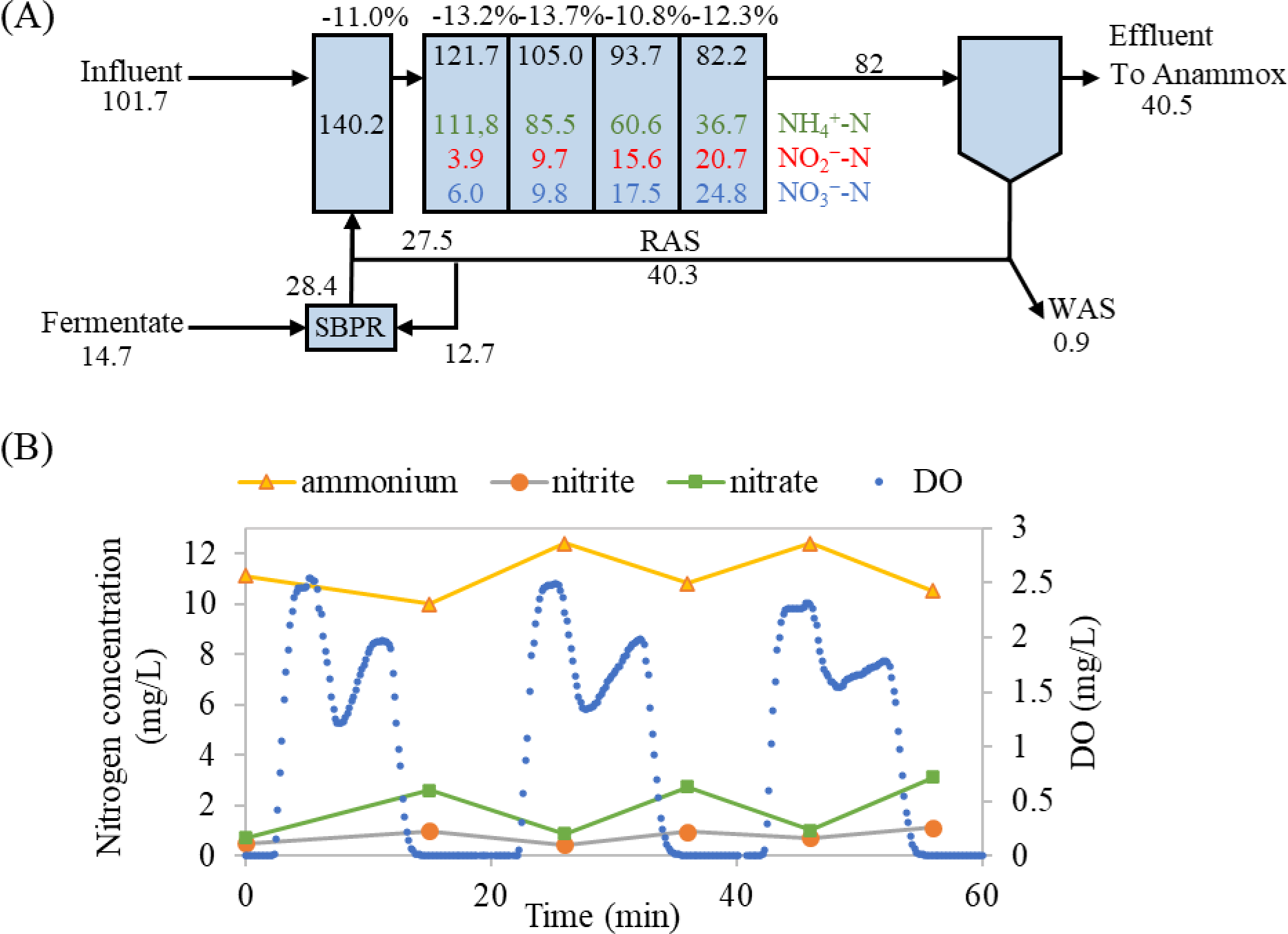
(A) Nitrogen mass balance of the pilot plant in study period (units are in g/d; values with the percentage sigh represent the removal rate, values of TIN, ammonia, nitrite, and nitrate are colored with black, green, red, and blue, respectively The B-stage effluent will go into the following anammox reactor; (B) Intermittent aeration profile taken on Day 252, showing nitrite accumulating during aerobic periods and being denitrified during anoxic periods. Reproduced from McCullough et al. (2023).

### 3.2 Nitrite accumulation due to partial nitrification

Since nitrite can be accumulated either through partial nitrification (PN) or partial denitrification (PDN) pathways, both AOB/NOB activities were evaluated over long-term on site to identify the dominant mechanisms involved in the nitrite accumulation (Figure 1). Examination of the N species during the aerobic/anoxic cycle showed that nitrite only accumulated in aeration periods while it was consumed after aeration was off (Figure 2B), suggesting that nitrite accumulation was likely due to PN rather than PDN. This was further supported by the AOB and NOB activities assessment. During the study period, the average AOB and NOB activities were 1.9 ± 0.9 and 1.1 ± 1.0 mgN/gSS/h, respectively (Figure 1). AOB activities were higher than the NOB activities during the period of study, consistent with the nitrite accumulation via PN (Figure 1). This evidence led to our conclusion that the higher nitrite accumulation in this study period is likely attributed to, at least mostly, PN.

### 3.3 Unknown AOB that contributed to the High AOB activity

#### 3.3.1 Low and negligible known AOB, NOB abundance

Temporal Microbial communities via 16S rRNA gene sequencing analysis revealed the microbial community structure dynamics and patterns during the study period (Figure 3). The average relative abundance of known AOB, including *Nitrosomonas* and *Nitrosomonadaceae* were remarkably low at 0.07% and 0.2%, respectively. The only known NOB is *Nitrospira*, has a relative abundance ranging from 1.7-6%. Their abundances were depicted in Figure 1. to show the variation trends. In our system, the relative abundances of both known AOB and NOB are significantly lower than that typically reported in mainstream PN/A systems (0.23-4.4% and 0.2- 53%) (summarized in Table S2). Particularly interesting, the relative abundance of *Nitrospira* was much higher than the known AOB, which did not align with the higher ammonia oxidation activity compared to nitrite oxidation activity.

**Figure 3.**
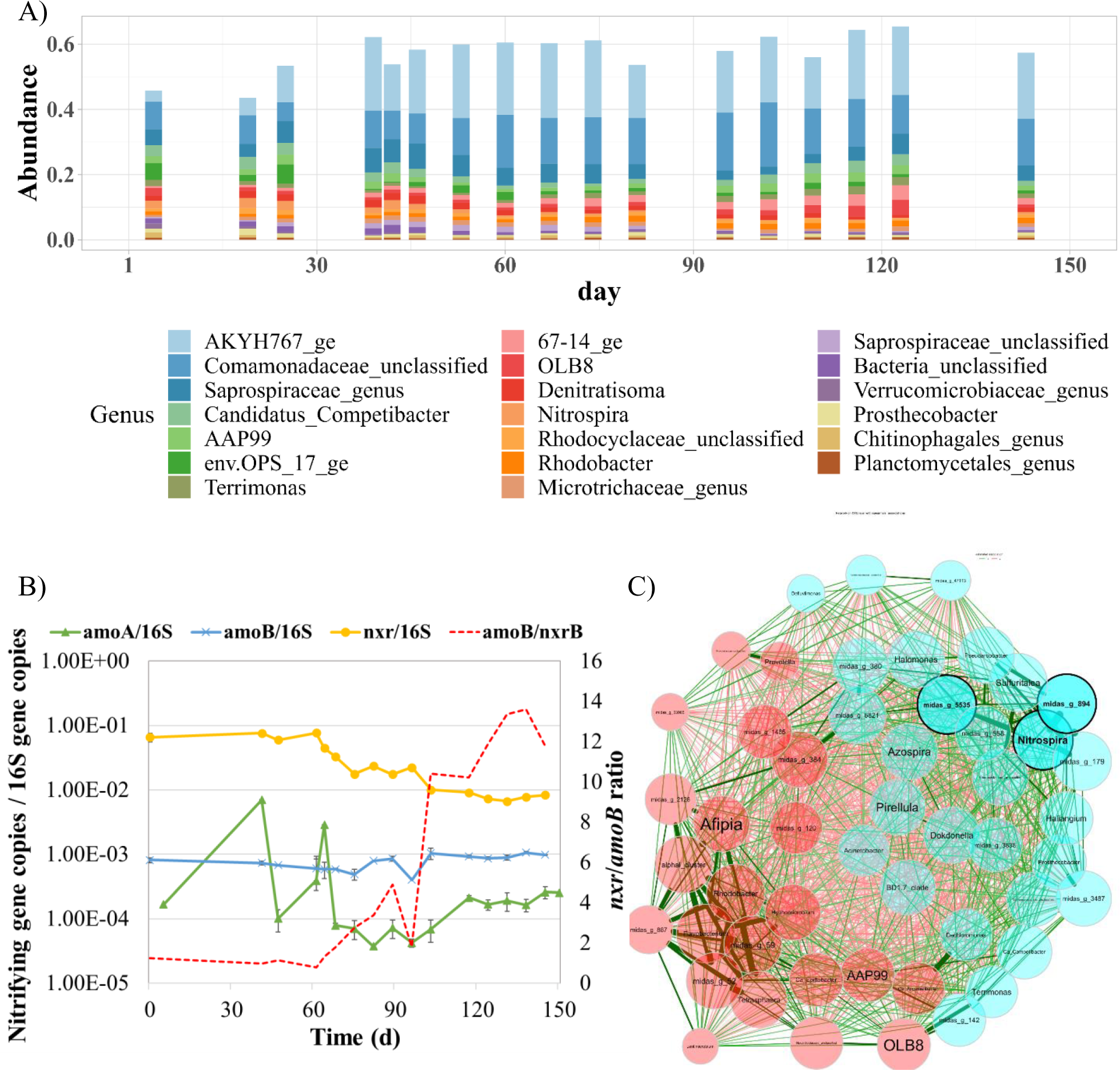
(A) Relative abundance of microbial community during nitrite accumulation period at genus level by 16S rRNA sequencing. (B) Quantitative nitrifying genes, *amoA* indicating ordinary proteobacteria AOB, *amoB* indicating comammox *Nitrospira*, and *nxr* indicating NOB *Nitrospira*, normalized by 16S rRNA genes. (C) Network analysis on OTU level with Spearman associations; The size of the nodes is proportional to its own total degree. The width of the edges is proportional to the correlation between the two nodes to which it corresponds. Positive and negative correlations between taxa (nodes) are indicated by blue and red color of the edges respectively. Detailed network analysis figure can be found in Figure S5.

To further verify the abundances of nitrifiers in the sludge, the *amoA* genes in AOB and the *nxr* genes in NOB were quantified and normalized to 16S rRNA genes (Figure 3B). The average *nxr*/16S ratio was 8.3−10^−3^ in this study period. Generally, *Nitrospira* genome harbor 1-6 copies of *nxr* (Palomo et al., 2018; Pester et al., 2014), while AOB in betaproteobacteria contains 2-3 copies of *amoA* (Norton et al., 2002). Considering the copy difference, the abundance of *nxrB* was still one order of magnitude higher than *amoA,* suggesting a higher known *NOB* abundance than known AOB abundance (Figure 3B). Both 16S sequencing and functional gene quantification confirmed that the abundance of known AOB (*Nitrosomonas* and *Nitrosomonadaceae*) was lower than NOB (*Nitrospira*) throughout the treatment periods, yet the system still achieved PN. In addition, the correlation analysis demonstrates that the microbial community changes have significant impact on the effluent nitrite accumulation (Figure S5). This result implies either the low abundance of nitrification communities played a significant role or some group of organisms carried out ammonia oxidation yet was unknown to us.

Theoretically, the numerical ratio of AOB to NOB in a balanced nitrifying system should be higher than 2:1 (Winkler et al., 2012). However, the current investigation shows a disparity between the rates of nitrite and ammonia oxidation, and the abundance of nitrifying biomass. Similar observations have been reported before (Wang et al., 2015; Fitzgerald et al., 2015; Zhang et al., 2018; Gao et al., 2021). Wang et al. (2015) observed NOB:AOB ratios from 9:1 to 5000:1 in five activated sludge reactors through qPCR. Zhang et al. (2018) achieved stable nitrite accumulation with higher abundance of *Nitrospira* comparing to AOB. Several scenarios can potentially explain the higher relative abundance of known NOB but lower nitrite oxidation activity in our system: 1) there might be a group of *Nitrospira* who can perform not only nitrite oxidation, but also ammonia oxidation, such as complete ammonia oxidizer (comammox); 2) underestimation of AOB in the system due to unknown AOB; and 3) some substances in wastewater could have more severe inhibitory effects on NOB than AOB.

#### 3.3.2 Comammox *Nitrospira* may contribute to nitrite accumulation

Roots et al., (2019) reported that the abundance of Comammox *Nitrospira* was much higher than both AOB and ammonia oxidizing archaea (AOA) in their in nitrification reactor (Roots et al., 2019), challenging the commonly held assumption that comammox play a minor role in wastewater treatment bioreactors. Among the versatile *Nitrospira* genus, the comammox *Nitrospira* encode a phylogenetically distinct *amo* gene from the canonical AOB and AOA, allowing them to fully oxidize ammonia to nitrate (Daims et al., 2016; Koch et al., 2015). High presence of comammox was reported in nitrite accumulating condition, indicating a potential correlation between the comammox and nitrite accumulation, given the higher ammonia oxidation rate than the nitrite oxidation rate in comammox *Nitrospira* (Shao and Wu, 2021). A recent isotope labelling test has proven that comammox species (*N. nitrosa* and *N. nitrificans*) provide nitrite for anammox bacteria (*Brocadia*) (van Kessel et al., 2015). The pure cultured *N. inopinata* was recorded to accumulate nitrite during complete nitrification (Kits et al., 2017). Given the function versatility of comammox species and the high abundance of *Nitrospira*, it’s possible that a portion of the comammox *Nitrospira* in our system functioned as AOB, leading to the accumulation of nitrite.

The presence of comammox *Nitrospira* in our system was determined by quantifying the *amoB* gene that was specifically found in comammox (Cotto et al., 2020; Roots et al., 2019) (Figure 3B). Results indicated an occurrence of higher abundance of comammox *Nitrospira* and nitrite accumulation. In addition, *amoB* was present and remained much more stable than both *nxrB* and *amoA* (Figure 3B). The average of *amoB* ratio was 7.8−10^−4^ per 16S gene copies. The *amoB*/*nxrB* ratio was deployed to indicate the relative abundance of comammox *Nitrospira* to the total *Nitrospira*. Results clearly indicate that comammox *Nitrospira* abundance increased from 1% to 13.6% during nitrite accumulation period. Considering that *Nitrospira* genome contains 1-5 times more copies of *nxr* (1-2 copies in most comammox genomes but 4 copies in *N. nitrificans* and 5 copis in *N. mosocoviensis*) than *amo* (single copy) (Palomo et al., 2018; Pester et al., 2014), the relative abundance of comammox *Nitrospira* to the total NOB ranges from 13.6% to 68% in our system.

The increasing abundance of comammox *Nitrospira* motivated us to employ the oligotyping analysis to further elucidate the micro-diversity of *Nitrospira*. This analysis identified 3 dominant oligotypes of *Nitrospira* and revealed their changes in microbial communities over time (Figure 4B).

**Figure 4.**
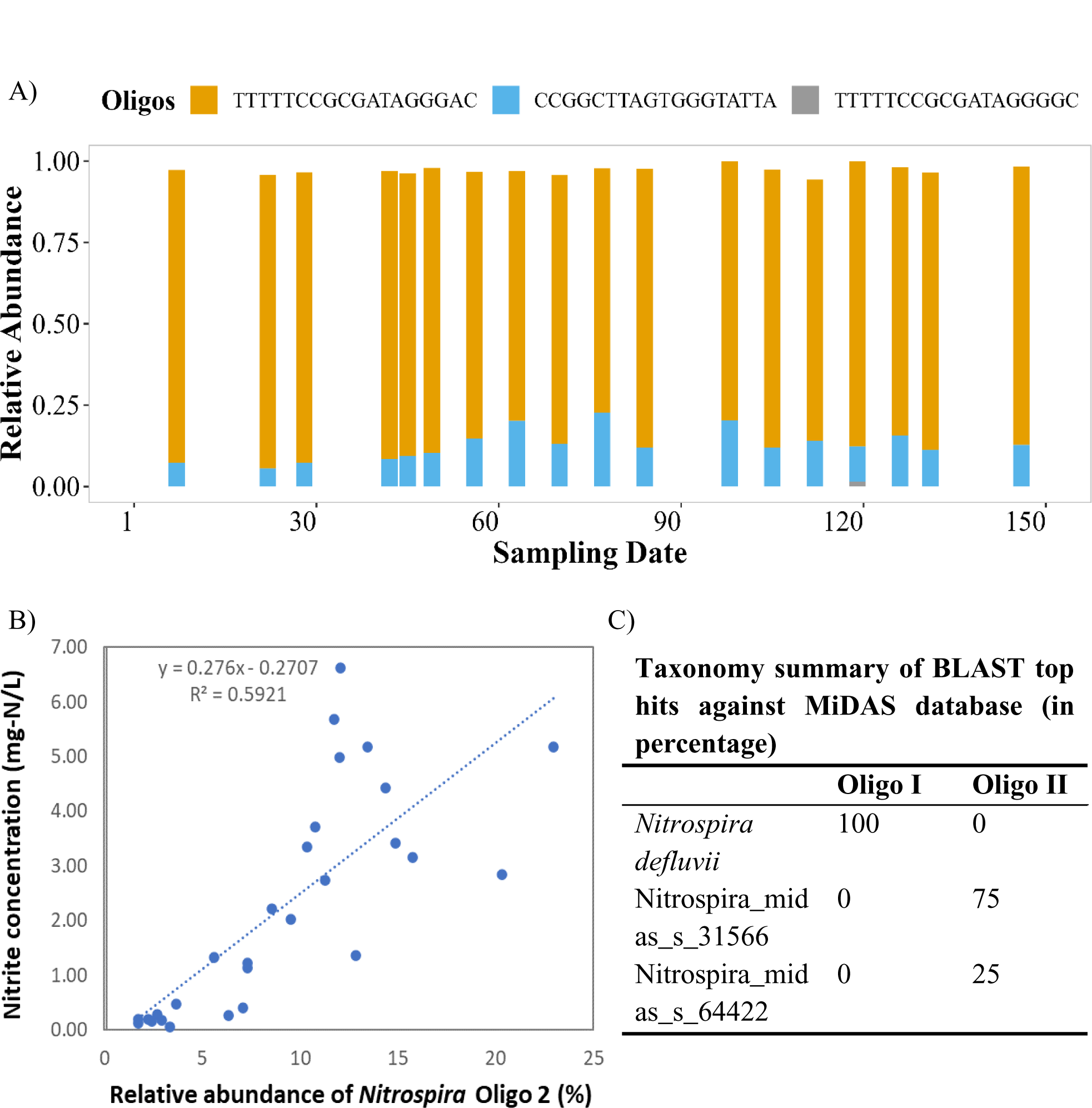
(A) Oligotypes (3 total) of 16S rRNA gene amplicon sequences classified as genus *Nitrospira* throughout the investigation period; (B) Positive correlation between the relative abundance of oligotype II (CCGGCTTAGTGGGTATTA) and the nitrite concentration; (C) Blast results of taxonomy identification of oligotypes I and II against MiDAS4 database.

Though the oligotype 1 consistently dominated the *Nitrospira* (> 71%), oligotype II gradually increased in tandem with the increasing nitrite accumulation. The similar increasing trends of *amoB*/*nxrB* and the relative abundance of oligotype II indicates a positive relationship between the comammox and the oligotype II *Nitrospira*. Note that comammox *Nitrospira* do not form a monophyletic group in 16S rRNA gene sequence or *nxrB*-based phylogenetic analyses, but are instead interspersed with strict nitrite-oxidizing *Nitrospira* (Daims et al., 2015; van Kessel et al., 2015). Different comammox *Nitrospira* encode *amo* orthologs that dissimilar to each other, with only ∼60% amino acids identity between members of the different clades (Daims et al., 2015), making it difficult to design a uniform comammox qPCR primers (Beach and Noguera, 2019; Cotto et al., 2020). The *amoB* primer set used in this study was specifically optimized for clade A comammox (Cotto et al., 2020). However, it is possible that clade B comammox also played a role in our S2EBPR-PN/A processes, but current primers did not capture the whole comammox community. Nevertheless, oligotyping and qPCR analysis indicated that a group of strict nitrite- oxidizing *Nitrospira* decreased, while another group of *Nitrospira*, which might belong to comammox and can oxidize ammonia increased, partially contributing to the nitrite accumulation (r^2^ = 0.60, Figure 4B). BLASTing the two major oligotypes of *Nitrospira* again MiDAS4 database, indicating that all sequences in oligotype I were identified as *Nitrospira defluvii* (Figure 4C). By summarizing the individual taxonomic identifications of all sequences in oligotype II subcluster, oligotype II is closer to Nitrospira_midas_s_31566 than Nitrospira_midas_s_64422. Considering the positive correlation of the nitrite accumulation and oligotype II of *Nitrospira*, it is possible that Nitrospira_midas_s_31566 is a candidate comammox *Nitrospira*.

#### 3.3.3 Atypical ammonia oxidation bacteria

Fitzgerald et al. (2015) had proposed that the abundance of ammonia oxidizing organisms may be underestimated, particularly in low DO condition (Fitzgerald et al., 2015). Despite the presence of complete nitrification to nitrate, no known AOB were identified in their low DO nitrification systems. To address this issue, they did pure culture and found 5 isolates that had not previously been recognized as AOB but can oxidize NH_4_ under both autotrophic and heterotrophic conditions. They proposed that genus *Pseudomonas* and *Rhodococcus*, and family Xanthomonadaceae and Sphingomonas are involved in heterotrophic nitrification under low-DO conditions.

To further explore the potential ammonia oxidizers in our samples, the functional potential of bacterial assemblages was predicted with PICRUSt2 in both enzymes and pathways. The dominated enzyme genes are related to carbohydrate metabolism (e.g., glycolysis/gluconeogesis metabolism), amino acid metabolism, DNA polymerase, and electron transport, which is conceivable that all those functions are vital for microorganisms. Then we specifically aimed at genes that can oxidize ammonia into nitrite. However, only three OTUs have the potential to hold the methane/ammonia monooxygenase subunits (AMO, EC:1.14.18.3 and 1.14.99.39) that oxidize ammonia into hydroxylamine, and they all belong to unclassified genus under order Rhodocyclaceae. Another 11 OTUs belonging to Comamonadaceae_unclassified have the potential to hold hydroxylamine oxidase (HAO, EC: 1.7.2.6) that can further oxidize hydroxylamine to nitrite (Table S3 and Figure S6). Indeed, AMO and HAO were purified from heterotrophic denitrifiers back in 1996 (Moir et al., 1996; Wehrfritz et al., 1996), and heterotrophic nitrification has drawn a lot attention nowadays. 134 isolates were identified at the genus/species level, and members from family Comamonadaceae and order Rhodocyclaceae were reported to be aerobic denitrifiers (Duan et al., 2022).

In addition, the co-occurrence microbial patterns during nitrite accumulation periods revealed that genus *Nitrospira*, midas_g_894, and midas_g_5535 are the main hubs for this specific microbiome structure (Figure 3C). The latter two belong to the family Saprospiraceae and Rhodocyclaceae, respectively, and the species in those families have all been shown to play a role in autotrophic/heterotrophic nitrification (Duan et al., 2022; Mehrani et al., 2022).

Though the relative abundance of OTUs that are predicted to hold *hao* only represents a small fraction of the entire Comamonadaceae_unclassified group (Figure S6), the prediction guides us to look into the function played by Comamonadaceae. Though most heterotrophic nitrifiers can oxidize ammonia to N_2_O or N_2_ by a full nitrification and denitrification pathway, some of them are not capable of aerobic denitrification and ends with nitrite (Duan et al., 2022). A test with enriched Comamonadaceae (∼96%) showed ammonium oxidation potential under aerobic conditions, with nitrite as the main product (Bao and Li, 2017), indicating the importance of Comamonadaceae in PN. Some research also indicates that they contribute to both nitrogen and phosphorus removal and can function as PAO (Ge et al., 2015). Our unique coupling of A-B-stage with S2EBPR created unprecedented selection forces and conditions for both P and N removal relevant organisms in this system, which would conceivably lead to enrichment of organisms with versatile functions different from those in conventional EBPR or PDN systems for main-stream treatment. In the current study, the relative abundance of Comamonadaceae family remained stable at unusually high relative abundance level. (13%) in the study period (Figure 3A) and they might contribute to nitrite accumulation.

It’s important to note that the predicted functional potential was based on the current database, which only includes the sequences of known AMO. Considering the fact that the *amo* genes in comammox *Nitrospira* are phylogenetically distinct from previously identified AMOs (van Kessel et al., 2015), it is possible that some other groups of bacteria are oxidizing ammonia but cannot be captured by the current database.

### 3.4 Possible factors affecting nitrite accumulation

Another possible scenario to explain nitrite accumulation is that NOB activities were more severely inhibited than AOB. Multiple strategies have been postulated to suppress NOB in PN reactors, including low DO, transient anoxic condition, FNA inhibition, FA inhibition, and VFA composition (Eilersen et al., 1994; Jiang et al., 2019; Park et al., 2010; Wang et al., 2014). The FNA concentrations in both B-stage influent and the mixing zone (B-stage influent + flow from Side-stream reactor) are nearly zero, which did not provide inhibition pressure on NOB. The DO ranged between 0.2-1.8 mg/L in 4 intermittently aerated CSTRs, in line with the DO range that is suitable for PN (Cao et al., 2017; Regmi et al., 2014). However, more and more evidence show that DO regulation is not reliable to suppress NOB alone (Park et al., 2010).

Compared to the conventional B-stage nutrient removal system, the coupling of S2EBPR with PN created unique environmental selection conditions differ from those in conventional EBPR or short-cut N system without EBPR. A split of RAS will flow through S2EBPR, where the unique feature of S2EBPR system exerted different condition for nitrifying populations since they were exposed to extended longer anaerobic time (HRT of anaerobic time is 3.65 hrs.) with lower ORP (−383 ± 17 mv) in comparison to conventional nutrient removal processes where AOB and NOB generally experience shorter anoxic conditions (1-2 hrs) with shallow ORP level (∼-150 mv). In addition, high level (126.7 ± 72.1 mg/L as COD) and more complex VFA (Figure 5B and Table S4, containing acetate, propionate, butyric, valeric acids) were added in side-stream reactor. Indeed, the sCOD concentrations into SBRP highly covariates with the nitrite accumulation in B-stage (Figure 5A). The sCOD mass in S2EBPR in this Period is 288 mg/L. When the carbon mass in S2EBRP decreased at the end of the study period, the nitrite accumulation also declined. This trend cannot be observed in B-stage reactor, indicating that side stream EBPR reactor is the place where NOB was inhibited. Spearman correlation analysis indicated that the VFA concentration into S2EBPR abundance is negatively correlated with the abundance of *Nitrospira* but positively correlated with the abundance of midas_g_887, which belongs to family Comamonadaceae (Figure 5E). This influence is more profound when the VFA mass flow into S2EBPR is high (> 90 g/d). The unique conditions in the S2EBPR, including the low ORP environment and the VFAs concentration and composition, may play roles for the nitrite accumulation and they are further discussed in the following sections.

**Figure 5.**
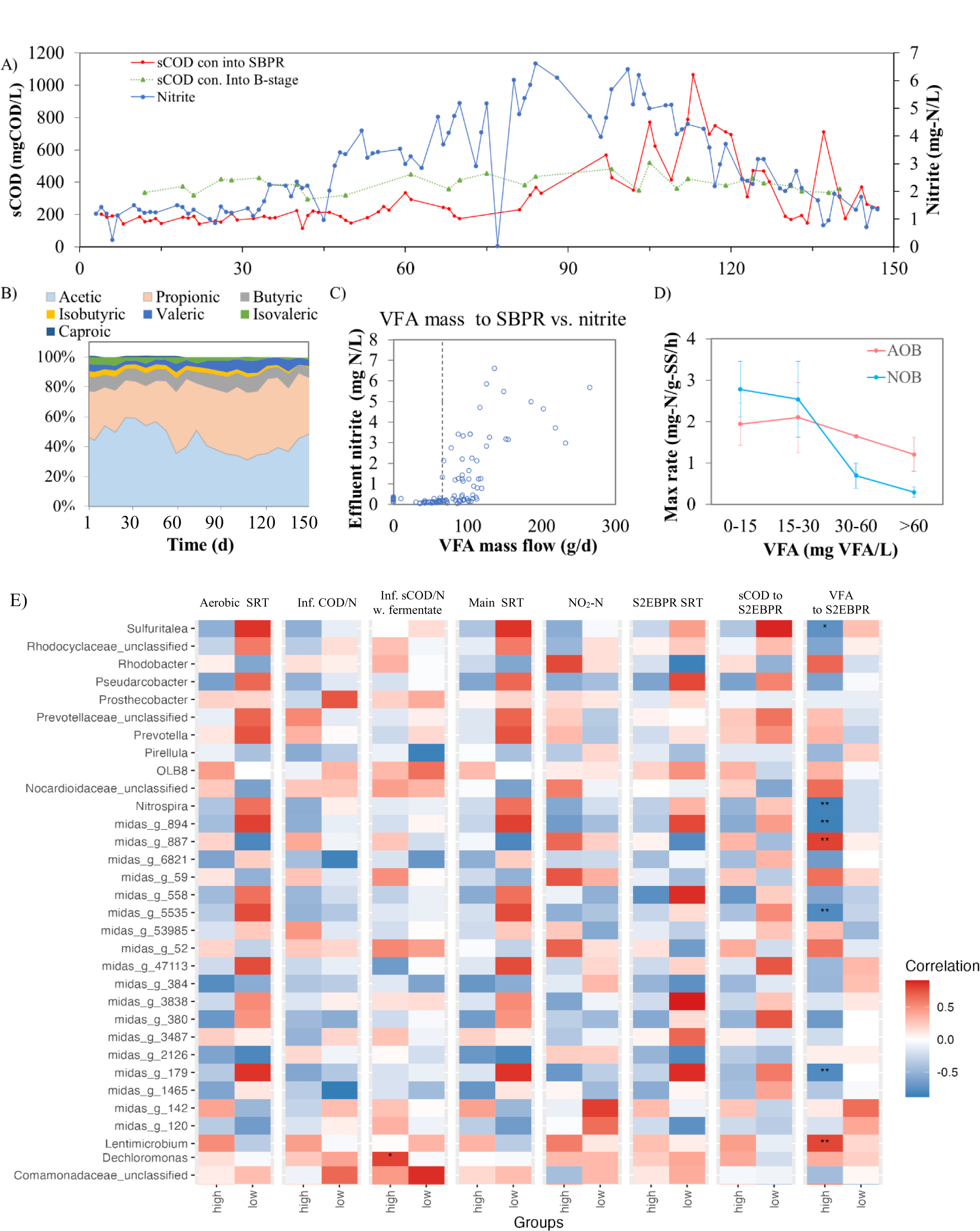
(A) Soluble COD concentrations in SBPR, B-stage and nitrite accumulation; (B) VFA speciation in A-stage fermentate; (C) VFA mass flow into SBPR and nitrite accumulation; (D) VFA concentration in SBPR and AOB/NOB activities; (E) The Spearman correlation between the most abundant 50 taxa and numerical environmental variables. The Spearman correlation test is performed and associated p-values are adjusted for multiple testing.

#### 3.4.1 VFAs Inhibit NOB activity and alter NOB metabolic pathway

In 1994, Eilersen et al. reported that nitrite oxidation is more easily to be inhibited by VFA than ammonia oxidation, especially acetic, propionic, and butyric acids (Table S5). Especially, the butyric acid exerted strong inhibition on NOB activity (IC_50_ = 399 mg/L) comparing to acetic (IC_50_ = 3707 mg/L) and propionic (IC_50_ = 3082 mg/L). Following research confirmed that NOB activity was more susceptible to VFA inhibition comparing to AOB and the inhibition level is proportional to the VFA molecular weight (Delgado et al., 2004; Gomez et al., 2000). All those researchers came to this conclusion since the nitrate production decreased when they added VFAs into NOB consortia. However, a later study showed that VFA (1500 mg/L) addition did not influence the CO_2_ fixation that is proportional to nitrite oxidation in NOB consortia (*Nitrospira* spp.), indicating other metabolic pathway was activated by VFAs rather than activity inhibition (Oguz et al., 2006). Though some *Nitrobacter* and *Nitrospira* benefit from the presence of simple organic compounds, e.g., pyruvate, formate, urea (Daims et al., 2016; Palomo et al., 2018), only comammox *Nitrospira* was reported to utilize fatty acids (acetate and propionate) (Xu et al., 2022), and even have potential to synthesize and degrade poly(3- hydroxybutyrate) (Palomo et al., 2018; Yang et al., 2020).

Comparing to the conventional WWTP that rely on mono external carbon source, such as methanol or acetate, our system’s A-stage fermentate contained a higher and stable proportion (58.9%) of high molecular size VFAs, including propionic, butyric, iso-butyric and isovaleric acids (Figure 5B). A-stage fermentate addition in S2EBPR may alter comammox *Nitrospira* towards organic matter utilization pathway and down regulate the nitrite oxidation enzymes. Notably, nitrite accumulation was not observed until the VFA mass flow was increased higher than 66 g/d (Figure 5C). The activity tests also indicated that higher residual VFA concentration (> 30 mg/L) in B-stage resulted lower NOB activity (in terms of nitrate production rates) (*p* < 0.01, Figure 5D), supporting the proposed role of residual VFAs in alternating the NOB activity (Figure 5E). All those results indicated that implementation of S2EBPR with VFAs supplement seem to favor the proliferation of versatile *Nitrospira* over canonical *Nitrospira*. This speculation was supported by quick increase of the relative abundance of comammox *Nitrospira* over the total *Nitrospira* community (indicated as *amoB*/*nxrB*) (Figure 4), although underlying mechanisms need further investigation.

#### 3.4.2 Longer exposure to anaerobic conditions with lower ORP impacts NOB

In our system, S2EBPR induced an extended anaerobic phase (∼3.7 hours) where AOB and NOB have to survive/maintain their activities for a longer time than typical duration in conventional WWTPs. Some research reported that the endogenous decay rate of AOB (0.020 d^−1^) was slower than that of NOB (0.035 d^−1^) in anoxic condition (Cui et al., 2019), this decay difference was also supported by the FISH staining analysis Morgenroth et al., (2000). However, other results showed identical decay rates of AOB and NOB under both anoxic and anaerobic conditions (Munz et al., 2011; Salem et al., 2006), The differences may have been due to wastewater characteristics, operation parameters, and the composition of the microbial communities and the related survival strategies (Geets et al., 2006; Hao et al., 2010). The ORP in S2EBPR was −383 ± 17 mV, which is significantly below the +100 to +350 mV in normal nitrification tank, and +50 to −50 mV in denitrification tank (Environmental, 2008). Generally, the endogenous decay rates of nitrifiers decreases with the decreasing ORP condition, as *b*_aerobic_ >> *b*_anoxic_ > *b*_anaerobic_ (Geets et al., 2006; Munz et al., 2011; Ruiz-Martínez et al., 2018; Salem et al., 2006). We assume that the actual decay rates in our system would be even lower than 0.02 d^−1^. Especially, the abundance of *Nitrospira* was much higher than AOB throughout the experiment periods, indicating that the decay differences between AOB/NOB did not play a significant role in NOB out-selection.

Other than low ORP, NOB is prone to be sensitive to the aerobic/anoxic perturbation. The nitrite oxidation enzyme reactivation time of NOB is longer than AOB when they are exposed to aerobic condition again (Gilbert et al., 2014; Kornaros et al., 2010). This characteristic has been applied as intermittent aeration strategy in PN tank to suppress the NOB activity (Kornaros et al., 2010). With more RAS split into S2EBPR (Figure S2), more AOB and NOB population would undergo anaerobic and VFA exposure. With less versability, AOB may slow down their metabolic activity while NOB may seek for other alternative pathway to maintain their energy.

This explains why they need a longer (enzyme) reactivation time (lag phase) to change back to autotrophic nitrite oxidation in intermittent aeration systems (Gilbert et al., 2014) and thus lead to nitrite accumulation.

## 4 Conclusions

Nitrite accumulation was observed along with phosphorus removal in a pilot-scale A/B-shortcut N-S2EBPR process treating municipal wastewater.

1. Nitrite accumulation was achieved by PN in A/B-shortcut N-S2EBPR system without canonical NOB out-selection, which was the higher ammonia oxidation activity than the nitrite oxidation rate.
2. *Nitrospira* dominated and was the center hub of the autotrophic nitrification community throughout the operation periods with one to two orders of magnitude higher than AOB, as indicated by 16s rDNA phylogenetic analysis and network analysis.
3. The contribution of comammox *Nitrospira* oligotype II (Nitrospira_midas_s_31566) in PN was suggested by the increased *amoB*/*nxr* ratio, the significant positive correlation between the abundance of *Nitrospira* Oligotype II and the ammonia oxidation activity.
4. The integration of S2EBPR into the A/B-shortcut N-S2EBPR seemed to exert distinct selective pressure on nitrifying community from those in conventional BNR systems.
5. Phylogenetic investigation suggested that ammonia oxidation enzymes (AMO and HAO) were embedded in heterotrophic organisms that belong to the family Comamonadaceae and order Rhodocyclaceae. These organisms have potential to function as heterotrophic nitrifiers in response to elevated VFA addition.

## Acknowledgement

This study was funded by the Water Research Foundation (WRF, Grant No. 4901). We gratefully acknowledge the financial support and scientific input from the WRF project team, particularly from Jim McQuarrie, Issac Avila, Dan Freedman, Rudy Maltos, Blair Wisdom (Denver Metro Wastewater Reclamation District), Beverley M. Stinson (AECOM), Christine deBarbadillo, Haydee De Clippeleir (District of Columbia Water and Sewer Authority), Paul Dombrowski (Woodard & Curran), Daniel R. Dair and Chandler H. Johnson (World Water Works) and James Barnard (Black & Veatch). Special thanks are given to the generous support of operators and staff of (Hampton Roads Sanitation District) for their assistance with sample and data collection.

