## Supplementary material for "*Comammox* and Unknown Candidate AOB Contribute to Nitrite Accumulation in an Integrated A-B stage process that Incorporates Side-stream EBPR (S2EBPR)": Suplemental Information

**Content**:


Supplementary Text 3

1. Figures

Figure S1. Schematic diagram of pilot A-B process with side-stream Bio-P plant (Figure is adopted from McCullough et al., (2020)) 4

Figure S2 The HRT and SRT of S2EBPR tank, the split portion of RAS into S2EBPR, and the B-stage effluent OP concentration during operation periods in this study (the days are re-set from day 1 for this period of study). 5

Figure S3 Nitrogen mass balance of the pilot plant before the nitrite accumulation period (units are in g/d; values with the percentage sign represent the removal rate, values of TIN, ammonia, nitrite, and nitrate are colored with black, green, red, and blue, respectively. 6

Figure S4 The fuzzy set ordination result of microbial community data for environmental variable, effluent nitrite concentration, between high VFA level (High in red, >90 g/d) and low VFA mass flow (Low in blue) (< 90 g/d) from A-stage fermentate to the S2EBPR reactor, with coefficients and significance level in the bracket. 7

Figure S5 Network analysis on OTU level with Spearman associations. 8

Figure S6 Abundance of OTUs that may contain *amo*/*pmo* or *hao* predicted by PICRUST2. 9

2. Tables

Table S1 The primer sets sequence and the reaction conditions for quantitative PCR performed in this study 10

Table S2 Relative abundance of AOB and NOB in partial nitrification process treating mainstream wastewater. 11

Table S3 OTUs that may contain *amo*/*pmo* or *hao* predicted by PICRUST2 12

Table S4 Relative fractions of acetic, propionic, butyric, isobutyric, valeric, isovaleric, and caproic of total VFA in Period I and II 14

Table S5 Critical concentrations (mM) of VFAs for inhibition of AOB and NOB 15

### Supplementary Text

To overcome the shortage of influent VFA and enable bio-P removal in main-stream treatment line, a side-stream reactor that received a split of return activated sludge and had the option of receiving supplemental carbon from A-stage sludge fermenter to enrich for PAOs and then allows for P removal in B-stage reactors (Figure S1). The HRT, SRT and the RAS split ratio in S2EBPR were summarized in Figure S2. Samples were collected over 7 months. For the initial phase without supernatant supplement from A-stage sludge fermenter, average effluent P level and removal efficiency was 1.8 ± 0.9  and  77% , respectively, As the fermentate flow rate and RAS split mass flow into S2EBPR increased to elevate the C/P (Figure S2), the average effluent phosphorus decreased from 1.8 ± 1.0 to 0.7 ± 0.85 mg-P/L, with consequent increase in P removal efficiency from 77% to 92%. The detailed discussion regarding to phosphorus removal performance and microbial population can be found in 4.

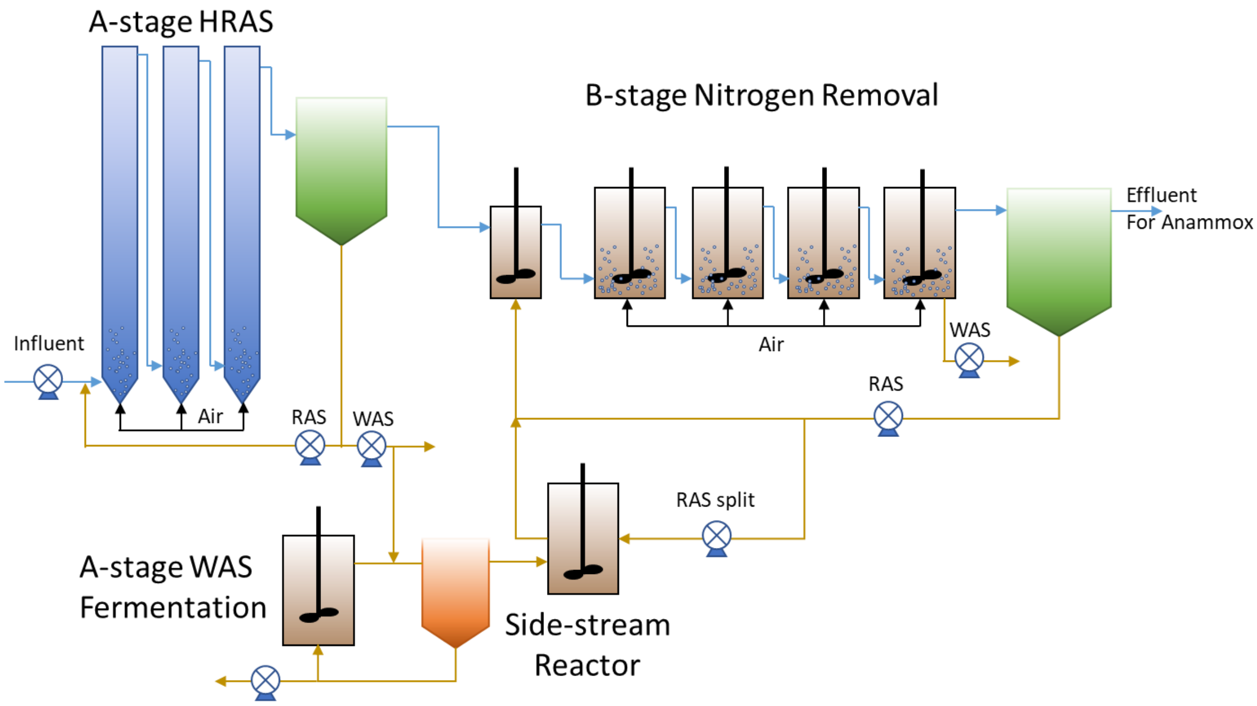


Figure S1. Schematic diagram of pilot A-B process with side-stream Bio-P plant (Figure is adopted from McCullough et al., (2020))

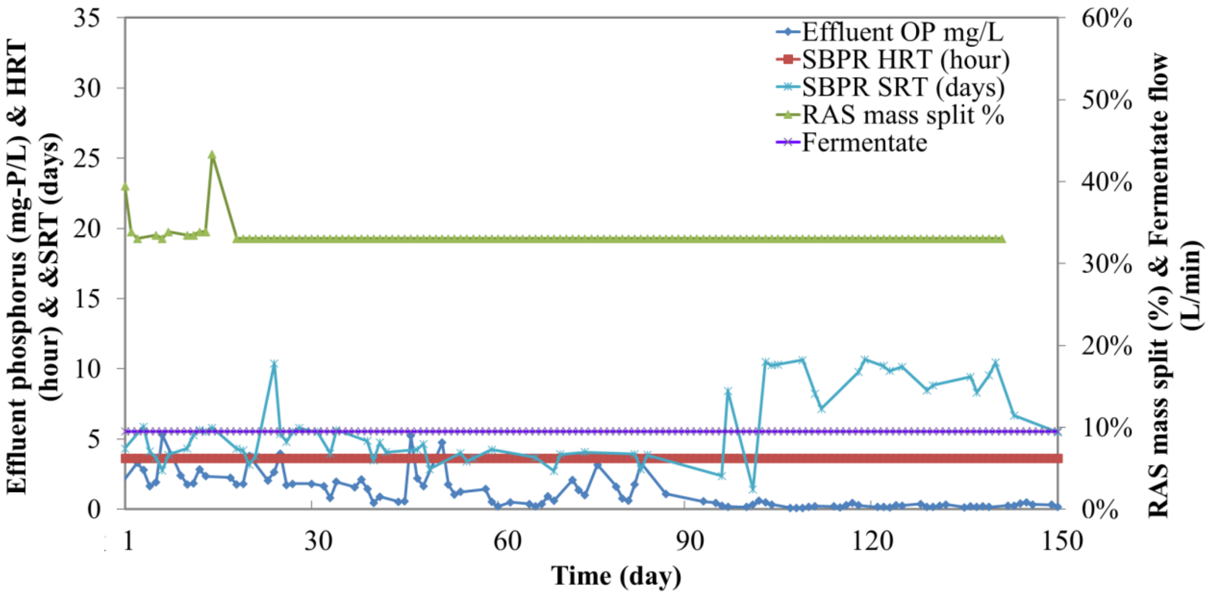


Figure S2 The HRT and SRT of S2EBPR tank, the split portion of RAS into S2EBPR, and the B-stage effluent OP concentration during operation periods in this study (the days are re-set from day 1 for this period of study).

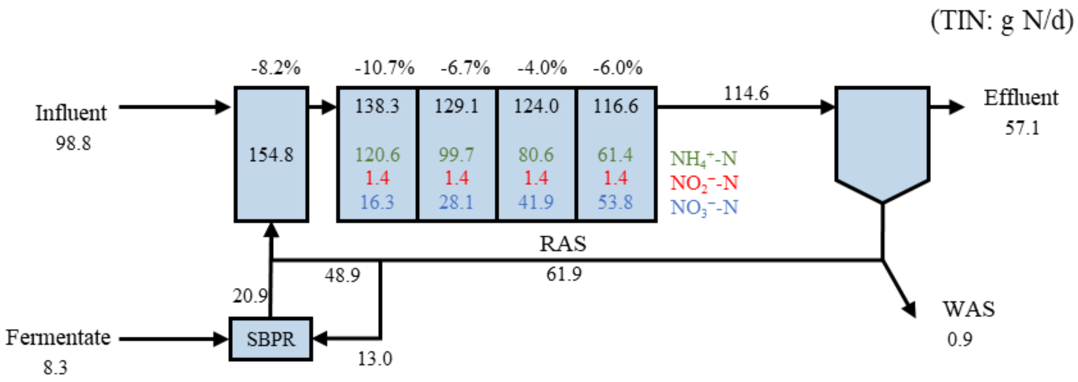


Figure S3 Nitrogen mass balance of the pilot plant before the nitrite accumulation period (units are in g/d; values with the percentage sign represent the removal rate, values of TIN, ammonia, nitrite, and nitrate are colored with black, green, red, and blue, respectively.

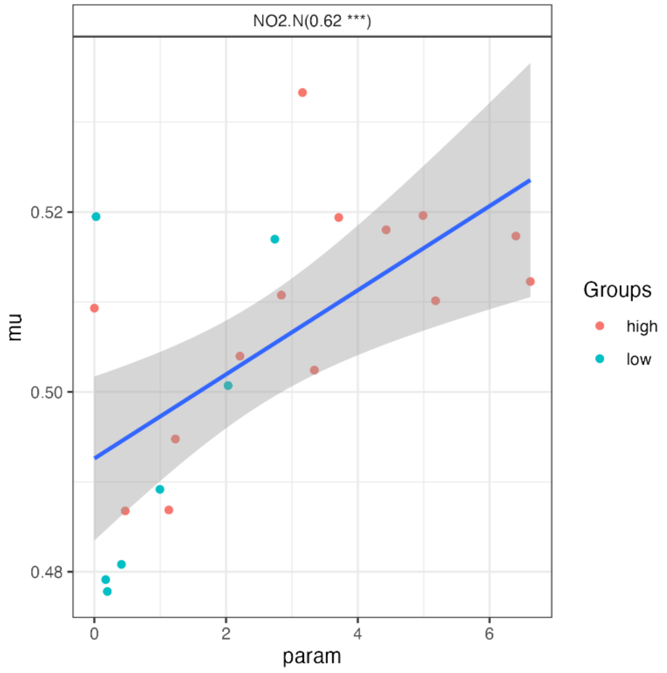


Figure S4 The fuzzy set ordination result of microbial community data for environmental variable, effluent nitrite concentration, between high VFA level (High in red, >90 g/d) and low VFA mass flow (Low in blue) (< 90 g/d) from A-stage fermentate to the S2EBPR reactor, with coefficients and significance level in the bracket. The less the coefficient is, the stronger the correlation between the changes in environment variable and the microbial abundance.

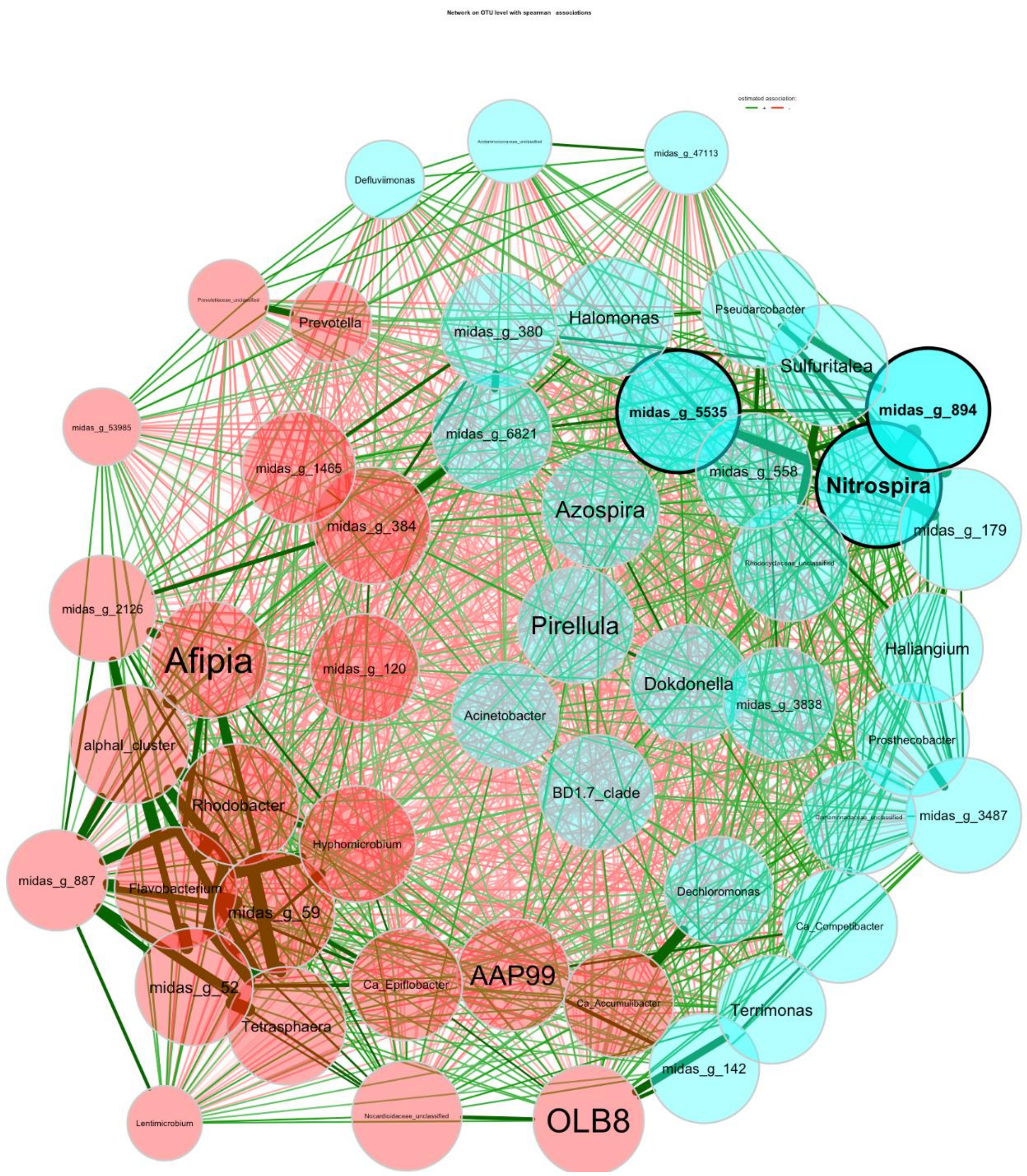


Figure S5 Network analysis on OTU level with Spearman associations; The size of the nodes is proportional to its own total degree. The width of the edges is proportional to the correlation between the two nodes to which it corresponds. Positive and negative correlations between taxa (nodes) are indicated by blue and red color of the edges respectively.

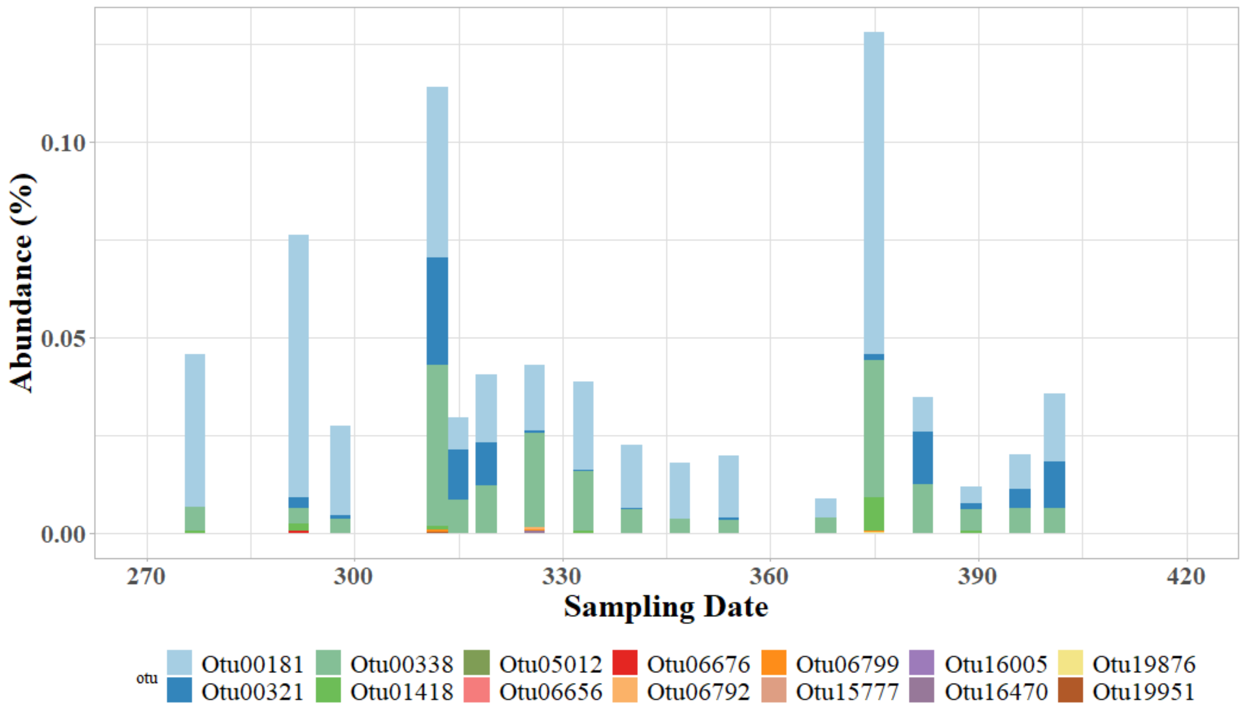


Figure S6 Abundance of OTUs that may contain *amo*/*pmo* or *hao* predicted by PICRUST2.

Table S1 The primer sets sequence and the reaction conditions for quantitative PCR performed in this study

| **Target** | **Primer** | **size (bp)** | **qPCR condition** | **Sequence** | **Reference** |
|---|---|---|---|---|---|
| Bacteria | 16S rRNA-341F/534R | 194 | 95°C for 3 minutes, then 40 cycles of 94°C for 30 seconds, 60°C for 45 seconds, and 72°C for 30 s. | 5'-CCTACGGGAGGCAGCAG/5'-ATTACCGCGGCTGCTGGCA | (He et al., 2007) |
| AOB | amoA-1F/2R | 491 | 95°C for 3 minutes, then 40 cycles of 95°C for 15 seconds, 54°C for 30 seconds, and 72°C for 1 minute. | 5'-GGGGTTTCTACTGGTGGT/5'-CCCCTCKGSAAAGCCTTCTTC | (Rotthauwe et al., 1997) |
| NOB | nxrB-169F/638R | 485 | 95°C for 5 minutes, then 35 cycles of 95°C for 40 seconds, 56.2°C for 40 seconds, and 72°C for 90 seconds. | 5'-TACATGTGGTGGAACA/5'-CGGTTCTGGTCRATCA | (Pester et al., 2014) |
| Comammox | amoB-148F/485R | 337 | 95°C for 3 minutes, then 40 cycles of 95°C for 15 seconds, 52°C for 45 seconds, and 72°C for 1 minute. | 5'-TGGTAYGAYACNGAATGGG/5'-CCCGTGATRTCCATCCA | (Cotto et al., 2020) |

Table S2 Relative abundance of AOB and NOB in partial nitrification process treating mainstream wastewater.

| **Ammonia con. mg-N/L** | **Configuration** | **AOB** | **NOB** | **Reference** |
|---|---|---|---|---|
| 25 ± 7 | Bench-scale Plug-flow reactor | Nitrosomonas, f_Nitrosomonadaceae 0.4-0.23% [a] | Nitrospira 1.07-0.20% [a] | (Feng et al., 2021) |
| 47.1 ± 4.4 | Bench- and pilot-scale Bottom-feed SBR | Nitrosomonadaceae 4.3-0.3% [a] | Nitrotoga 2.3-0.8% [a] | (Hausherr et al., 2022) |
| 14.3 ± 4.7 | Bench-scale SBR | Nitrosomonas 4.4%-0.8% [a] | *Nitrospira* 3.1%-53% [a] | (Roots et al., 2019) |
| 58.1 | Bench-scale SBR | β-AOB 3-4%[b] | *Nitrospira* *Nitrobacter* 2.5-4% [b] | (Guo et al., 2010) |
| 19.8-49.9 | Bench-scale AO-SBR | *Nitrosomonas* 0.01-0.09 [a] | *Nitrospira* 0.5-0.78 [a] | (Feng et al., 2022) |
| 27.57-50 | A2O, Oxidation ditch, MAO* | *Nitrosomonas* 1.27% [a] | *Nitrospira* 4.02% [a] | (Yao and Peng, 2017) |
| 44.5-72.6 | Bench-scale AO-SBR | *Nitrosomonas and Nitrosomonadaceae* 1.15-1.46%[a] | *Nitrospira* 1.34-1.17%[a] | (Cui et al., 2019) |

AO-SBR denotes the SBR is operated under anaerobic/aerobic cycle mode;

A2O denotes anaerobic/anoxic/aerobic process

MAO denotes multiple AO process

* the research investigated 10 full-scale WWTPs receiving both domestic and industrial wastewater with full nitrification and denitrification process

[a] the abundance was measured by 16S sequencing;

[b] the abundance was measured by fluorescence *in situ* hybridization.

Table S3 OTUs that may contain *amo*/*pmo* or *hao* predicted by PICRUST2

| **OTU** | **Taxon** |
|---|---|
| OTU00181 | Bacteria(100);Proteobacteria(100);Gammaproteobacteria(100);Burkholderiales(100);Comamonadaceae(67);Comamonadaceae_unclassified(94) |
| OTU00321 | Bacteria(100);Proteobacteria(100);Gammaproteobacteria(100);Burkholderiales(100);Comamonadaceae(67);Comamonadaceae_unclassified(100) |
| OTU00338 | Bacteria(100);Proteobacteria(100);Gammaproteobacteria(100);Burkholderiales(100);Comamonadaceae(67);Comamonadaceae_unclassified(99) |
| OTU01418 | Bacteria(100);Proteobacteria(100);Gammaproteobacteria(100);Burkholderiales(100);Comamonadaceae(67);Comamonadaceae_unclassified(99) |
| OTU05012 | Bacteria(100);Proteobacteria(100);Gammaproteobacteria(100);Burkholderiales(100);Comamonadaceae(67);Comamonadaceae_unclassified(80) |
| OTU06656 | Bacteria(100);Proteobacteria(100);Gammaproteobacteria(100);Burkholderiales(100);Comamonadaceae(67);Comamonadaceae_unclassified(67) |
| OTU06676 | Bacteria(100);Proteobacteria(100);Gammaproteobacteria(100);Burkholderiales(100);Rhodocyclaceae(100);Rhodocyclaceae_unclassified(100) |
| OTU06792 | Bacteria(100);Proteobacteria(100);Gammaproteobacteria(100);Burkholderiales(100);Comamonadaceae(67);Comamonadaceae_unclassified(67) |
| OTU06799 | Bacteria(100);Proteobacteria(100);Gammaproteobacteria(100);Burkholderiales(100);Comamonadaceae(67);Comamonadaceae_unclassified(67) |
| OTU15777 | Bacteria(100);Proteobacteria(100);Gammaproteobacteria(100);Burkholderiales(100);Rhodocyclaceae(100);Rhodocyclaceae_unclassified(100) |
| OTU16005 | Bacteria(100);Proteobacteria(100);Gammaproteobacteria(100);Burkholderiales(100);Comamonadaceae(67);Comamonadaceae_unclassified(100) |
| OTU16470 | Bacteria(100);Proteobacteria(100);Gammaproteobacteria(100);Burkholderiales(100);Comamonadaceae(67);Comamonadaceae_unclassified(100) |

| | |
|---|---|
| OTU19876 | Bacteria(100);Proteobacteria(100);Gammaproteobacteria(100);Burkholderiales(100);Comamonadaceae(67);Comamonadaceae_unclassified(100) |
| OTU19951 | Bacteria(100);Proteobacteria(100);Gammaproteobacteria(100);Burkholderiales(100);Rhodocyclaceae(100);Rhodocyclaceae_unclassified(100) |

Table S4 Relative fractions of acetic, propionic, butyric, isobutyric, valeric, isovaleric, and caproic of total VFA in Period I and II

| **Statistics** | **Acetic** | **Propionic** | **Butyric** | **Isobutyric** | **Valeric** | **Isovaleric** | **Caproic** |
|---|---|---|---|---|---|---|---|
| | **Period I** | | | | | | |
| **Mean** | 38.7% | 34.7% | 11.2% | 3.3% | 8.2% | 3.6% | **0.5%** |
| **Standard Error** | 1.0% | 1.3% | 0.6% | 0.2% | 0.4% | 0.2% | **0.1%** |
| **Median** | 38.5% | 34.0% | 10.4% | 3.1% | 7.9% | 3.6% | **0.7%** |
| **Standard Deviation** | 5.8% | 7.1% | 3.4% | 0.9% | 2.1% | 1.0% | **0.4%** |
| **Minimum** | 30.1% | 22.6% | 6.1% | 1.9% | 4.3% | 1.5% | **0.0%** |
| **Maximum** | 54.4% | 49.8% | 16.7% | 5.2% | 12.4% | 6.2% | **1.1%** |
| | **Period II** | | | | | | |
| **Mean** | 44.7% | 36.8% | 8.3% | 2.6% | 4.9% | 2.6% | **0.4%** |
| **Standard Error** | 1.9% | 1.8% | 0.3% | 0.2% | 0.4% | 0.4% | **0.1%** |
| **Median** | 44.8% | 38.6% | 8.5% | 2.7% | 4.7% | 2.3% | **0.2%** |
| **Standard Deviation** | 8.9% | 8.3% | 1.5% | 1.1% | 1.8% | 1.7% | **0.4%** |
| **Minimum** | 31.0% | 24.8% | 5.2% | 0.0% | 2.3% | 0.0% | **0.0%** |
| **Maximum** | 59.5% | 50.0% | 10.5% | 4.3% | 8.3% | 5.8% | **1.0%** |

Table S5 Critical concentrations (mM) of VFAs for inhibition of AOB and NOB

| VFAs | Ammonia oxidation | | | | | Nitrite oxidation | | | | |
|---|---|---|---|---|---|---|---|---|---|---|
| | $I_k$ (mM) | n (-) | $I_c$ (mM) | $I_c$ (mg/L as VFA) | $I_c$ (mg/L as COD) | $I_k$ (mM) | n (-) | $I_c$ (mM) | $I_c$ (mg/L as VFA) | $I_c$ (mg/L as COD) |
| Acetic | - | | | - | - | 115.39 | 1.00 | 57.70 | 3465 | 3707 |
| Propionic | - | | | - | - | 67.98 | 1.00 | 33.99 | 2041 | 3082 |
| n-Butyric acid | - | | | - | - | 32.83 | 0.17 | 3.65 | 219 | 399 |
| Isobutyric | 6.17 | 0.46 | 1.68 | 148 | 270 | 7.53 | 0.77 | 3.11 | 187 | 340 |
| Valeric | 37.21 | 0.36 | 8.22 | 839 | 1712 | 75.09 | 1.00 | 37.55 | 2255 | 4599 |
| Isovaleric | 6.29 | 0.32 | 1.25 | 128 | 261 | 7.20 | 0.51 | 2.14 | 129 | 263 |
| Caproic | 36.17 | 0.45 | 9.69 | 1126 | 2499 | 80.63 | 1.00 | 40.32 | 2421 | 5375 |

$I_K$ and n stand for critical concentration of inhibitors and the exponent n. They were fitted from regression model of critical concentration, which all activity ceases according to the empirical model: $V_I = V\left(1 - \frac{I}{I_K}\right)^n$ for $I \leq I_K$. Data collected from (Eilersen et al., 1994)).

$I_C$ denotes the half inhibition concentration of VFAs calculated based on the model. They are converted to COD for better comparison.
